# Insulin signaling regulates R2 retrotransposon expression to orchestrate transgenerational rDNA copy number maintenance

**DOI:** 10.1101/2024.02.28.582629

**Authors:** Jonathan O Nelson, Alyssa Slicko, Amelie A Raz, Yukiko M Yamashita

## Abstract

Preserving a large number of essential yet highly unstable ribosomal DNA (rDNA) repeats is critical for the germline to perpetuate the genome through generations. Spontaneous rDNA loss must be countered by rDNA copy number (CN) expansion. Germline rDNA CN expansion is best understood in *Drosophila melanogaster*, which relies on unequal sister chromatid exchange (USCE) initiated by DNA breaks at rDNA. The rDNA-specific retrotransposon R2 responsible for USCE-inducing DNA breaks is typically expressed only when rDNA CN is low to minimize the danger of DNA breaks; however, the underlying mechanism of R2 regulation remains unclear. Here we identify the insulin receptor (InR) as a major repressor of R2 expression, limiting unnecessary R2 activity. Through single-cell RNA sequencing we find that male germline stem cells (GSCs), the major cell type that undergoes rDNA CN expansion, have reduced InR expression when rDNA CN is low. Reduced InR activity in turn leads to R2 expression and CN expansion. We further find that dietary manipulation alters R2 expression and rDNA CN expansion activity. This work reveals that the insulin pathway integrates rDNA CN surveying with environmental sensing, revealing a potential mechanism by which diet exerts heritable changes to genomic content.

## Introduction

Ribosomal DNA (rDNA) loci are essential regions of the genome containing hundreds of tandemly repeated ribosomal RNA (rRNA) genes. The repetitive nature of rDNA loci makes them prone to undergo intrachromatid recombination between rDNA copies, leading to rDNA copy number (CN) reduction (Warmerdam and Wolthuis 2019) **(Fig 1A)**. Cell viability relies on a high number of rDNA repeats, thus protection against continual rDNA CN reduction is critical, especially in the lineages that continue through long time scale, such as the metazoan germline that passes the genome through generations (Nelson et al. 2019). In these cell types, the expansion of rDNA CN plays a critical role to counteract spontaneous rDNA CN reduction and achieve long term rDNA CN maintenance (Lu et al. 2018; Nelson et al. 2023b).

**Figure 1.**
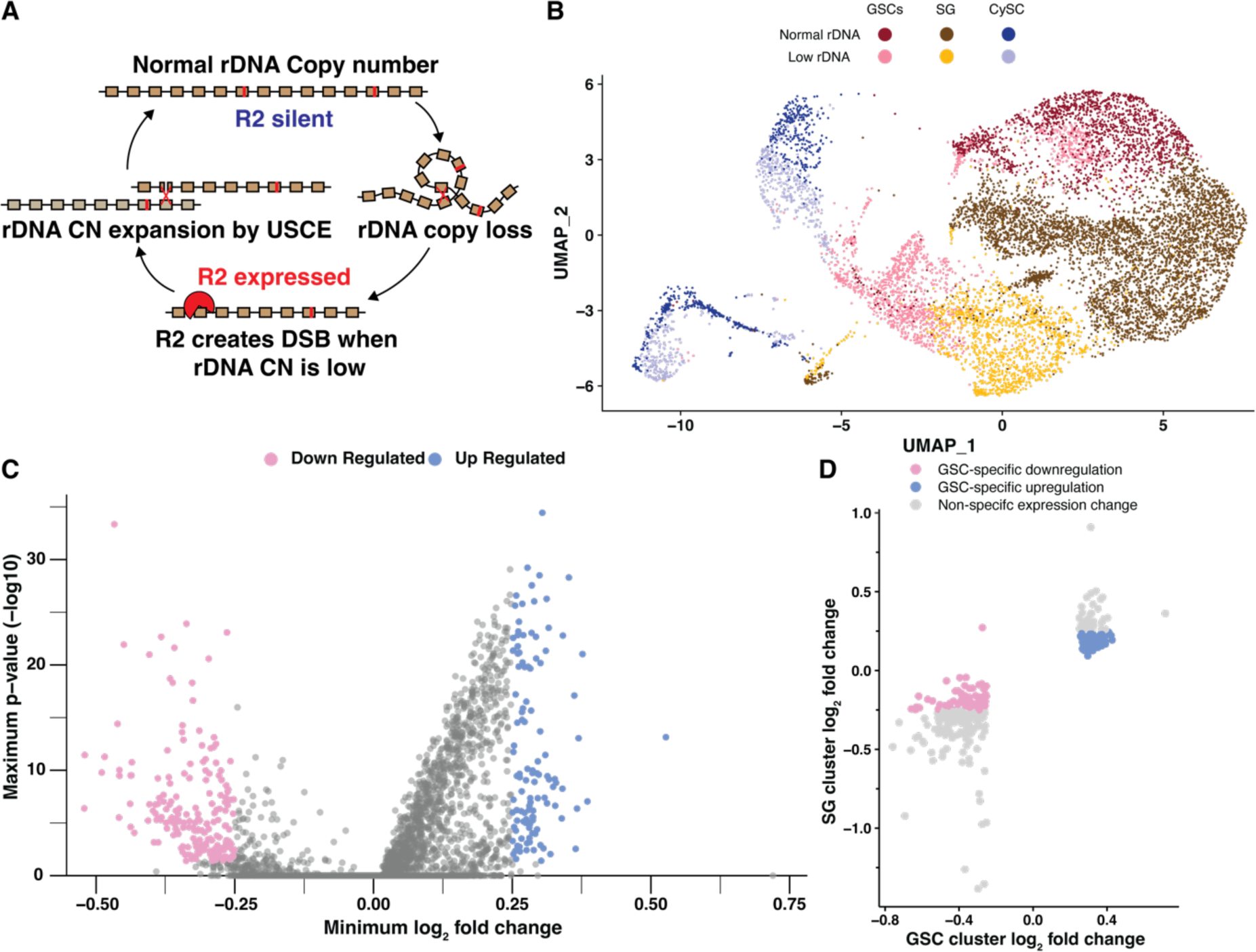
Single cell RNA sequencing reveals candidate repressors of rDNA magnification. **A** Model of R2 function in rDNA copy number (CN) maintenance. Expression of R2 when rDNA CN is reduced causes rDNA-specific R2 endonuclease activity to create double stranded DNA breaks (DSBs) at the rDNA locus, which can lead to rDNA CN expansion by unequal sister chromatid exchange (USCE) during their repair. **B** UMAP 2-dimensional reduction of early germ cells and somatic cyst cells from combined low and normal rDNA CN *upd* over-expression testes. **C** Differential gene expression in GSCs from combined analyses (see methods). Lowest fold change and highest p-value produced for either analysis is displayed. **D** Differential gene expression in low rDNA CN GSCs and spermatogonia (SG) determined by cluster-based cell selection. Significant gene expression change indicated by a log_2_ fold change > 0.25 or < −0.25.

*Drosophila melanogaster* and budding yeast have long served as excellent models to investigate rDNA CN expansion, with germline rDNA CN expansion first being described in Drosophila in the 1960s as the phenomenon called ‘rDNA magnification’ (Ritossa 1968; Tartof 1973). Drosophila that harbors unusually low rDNA CN exhibits visible phenotypes such as thin bristles and cuticle defects, collectively called the ‘bobbed’ (bb) phenotype (Ritossa et al. 1966). rDNA magnification describes the process of bobbed fathers to produce offspring that have reverted to wild-type cuticles due to rDNA CN expansion within the father’s germline (Ritossa 1968). Recent studies suggest that rDNA magnification is a manifestation of the rDNA CN expansion mechanism that maintains rDNA against spontaneous CN reduction in the germline (Lu et al. 2018; Nelson et al. 2023b). The easily scorable cuticle phenotype of bobbed flies and its reversion due to rDNA CN expansion have served as a powerful paradigm to study germline rDNA CN expansion (Hawley and Marcus 1989).

rDNA magnification is thought to occur by unequal sister chromatid exchange (USCE) (**Fig 1A**) (Tartof 1974; Hawley and Marcus 1989). USCE is triggered by DNA double-strand breaks (DSBs), which leads to homology-dependent recombinational repair between misaligned copies, allowing one sister chromatid to acquire rDNA copies from the other sister chromatid **(Fig 1A)** (Tartof 1974). We recently found that an rDNA-specific retrotransposon R2 is responsible for DSB formation during magnification **(Fig 1A)** (Nelson et al. 2023b). R2 was shown to be required to maintain rDNA copy number through generations (Nelson et al. 2023b), representing a striking example of host-transposable element (TE) mutualism, wherein active TEs benefit (or are required for the survival of) the host (Nelson et al. 2023b; Cosby et al. 2019). Whereas R2’s ability to create DSBs at rDNA loci is required for rDNA magnification (Nelson et al. 2023b), unnecessary R2 transposition and DSB formation at rDNA threaten the stability of rDNA loci (Eickbush and Eickbush 2015). Therefore, for R2 to be beneficial to the host, there must be a mechanism that regulates R2 activity according to the host’s needs, such as when rDNA CN is critically low or physiological demand for rDNA copies is increased.

To achieve effective rDNA magnification, USCE occurs in asymmetrically dividing germline stem cells (GSCs), such that the sister chromatid that gained rDNA CN can be selectively retained within the GSC (Watase et al. 2022; Nelson et al. 2023a). This biased inheritance in USCE products during asymmetric GSC divisions can effectively expand rDNA CN through repeated rounds of USCE over successive GSC divisions (Nelson et al. 2023a). Additionally, R2 expression during rDNA magnification is restricted to GSCs, whereas differentiating germ cells (spermatogonia) do not express R2 to avoid DSBs in these cells that are particularly sensitive to DNA damage (Nelson et al. 2023a; Lu and Yamashita 2017). Therefore, GSCs appear to be uniquely capable of regulating R2 expression in response to rDNA CN to control the activity of rDNA magnification. It remains unknown how these cells sense the need for rDNA CN expansion and how such information leads to R2 derepression.

Here, using single cell RNA sequencing (scRNA-seq), we identified genes that are differentially regulated in GSCs in response to low rDNA CN. Through this analysis, we identified Insulin-like Receptor (InR) as a gene downregulated in GSCs under low rDNA CN. We found that InR is a negative regulator of R2 expression, thereby preventing unnecessary R2 expression when cells have sufficient rDNA CN. Further, we showed that the mechanistic target of rapamycin (mTor), a major effector of insulin/IGF signaling (IIS), also represses R2 expression. We propose that IIS transduces rDNA CN surveillance within GSCs to the regulation of rDNA magnification activity via its ability to repress R2 expression.

## RESULTS

### Single cell RNA sequencing identifies differentially expressed GSC genes upon rDNA CN reduction

We recently showed that rDNA magnification primarily occurs in GSCs and not the more differentiated germ cells, e.g. spermatogonia (SGs) (Nelson et al. 2023a). Accordingly, genes that regulate rDNA magnification likely have altered activity or expression within GSCs when rDNA CN is low. Thus, we sought to identify genes that are differentially expressed in GSCs under low rDNA CN (magnifying) vs. normal rDNA CN conditions. Drosophila rDNA loci reside on the sex chromosomes (X and Y), and low rDNA CN conditions can be generated by combining rDNA-deficient sex chromosomes (Ritossa et al. 1966; Ritossa 1968). We used *bb^Z9^ / Ybb^0^* males, which contain reduced X chromosome rDNA CN (*bb^Z9^*) and no Y chromosome rDNA (*Ybb^0^*), for ‘low rDNA CN’ conditions that induce rDNA magnification (Nelson et al. 2023b). As a ‘normal rDNA CN’ control, we used *bb^Z9^ / Ybb^+^* males, which contain sufficient rDNA CN on the Y chromosome, and are as genetically matched to *bb^Z9^ / Ybb^0^* males as possible while not inducing rDNA magnification (Nelson et al. 2023b). Each testis contains only ∼8-10 GSCs, as opposed to hundreds of more differentiated SGs, spermatocytes, and spermatids **(Fig S1A left)**, therefore making it challenging to isolate enough GSCs to identify GSC-specific transcript changes even using single cell RNA sequencing (scRNA-seq) (Li et al. 2022; Raz et al. 2023). To circumvent this challenge, we conducted scRNA-seq in animals with low or normal rDNA CN using testes overexpressing *upd* in their early germline (*nos>upd*). Upd expression in the early germline leads to overproliferation of GSCs, enriching our cell type of interest **(Fig S1A right)** (Kiger et al. 2001; Tulina and Matunis 2001). Importantly, we confirmed that *upd* overexpression does not interfere with the induction of *R2* expression in GSCs in response to low rDNA CN **(Fig S1B-D)**.

We sought to discover potential regulators of rDNA magnification through identifying genes with altered expression in GSCs with low rDNA CN. To do so, we analyzed the transcriptome of a total of 20,138 quality-verified cells from low and normal rDNA CN *upd*-expressing testes. Cell identity was initially assigned based on rDNA CN condition and previously-determined testis cell type expression signatures, revealing cells among all spermatogonial stages in our samples **(Fig S2A-B**, see methods) (Li et al. 2022; Raz et al. 2023). In order to enrich for true positives of differentially-expressed GSC genes, we used two complementary methods to select GSCs for gene expression analysis. In one method, we used iterative sub-clustering to isolate true GSCs, while excluding SG and suspected artifactual GSC:non-GSC cell doublets **(Fig 1C**, see methods). The advantage of this cluster-based method is that it included cells with transcriptional signatures similar to GSCs even if they did not express all of the specific markers used to distinguish GSCs. This method resulted in 3087 GSCs to use for differential expression analysis, which identified 721 significantly downregulated and 320 significantly upregulated genes in GSCs with low rDNA CN compared to GSCs with normal rDNA CN **(Fig S2C, Table S1)**. The other method identified GSCs based on the expression of specific markers within each cell, regardless of their assigned cluster (**Fig S2D,** see methods). The advantage of this expression-based method is that it included all cells that may be considered GSCs even if their overall expression profile failed to cluster with other GSCs. This second method produced 746 GSCs for differential expression analysis, and identified 247 significantly downregulated and 167 significantly upregulated genes in GSCs with low rDNA CN **(Fig S2E, Table S1)**. Selecting the overlap between these two analyses yielded 202 genes that are downregulated and 117 genes that are upregulated in low rDNA CN conditions in GSCs from both analyses **(Fig 1D, Fig S2F, Table S1)**. To further narrow down these candidates, we reasoned that regulators of rDNA magnification would have altered expression specifically in GSCs but not SG, since rDNA magnification is a unique feature of GSCs and not the more differentiated SG (Nelson et al. 2023a). Therefore, we assessed SG gene expression for all 319 differentially expressed genes in GSCs, and found that 71 downregulated and 84 upregulated GSC genes have no (or opposite) expression change in low rDNA CN SG, suggesting these genes change expression only in GSCs in response to rDNA CN **(Fig 1D, Table S2)**.

### rDNA magnification is repressed by insulin-like receptor (InR)

In this study, we focused on candidate negative regulators of rDNA magnification because their functional importance can be assayed relatively easily. We have previously shown that exogenous expression of R2 is sufficient to induce ‘ectopic’ rDNA magnification (Nelson et al. 2023b), and surmised that RNAi-mediated knockdown of negative regulators of rDNA magnification would have a similar effect. Ectopic rDNA magnification is observed in *bb^Z9^ / Ybb^+^*males, which do not normally expand rDNA CN at the *bb^Z9^* locus because they have sufficient rDNA CN. Ectopic rDNA magnification in these males can be observed by mating them to females harboring the rDNA deletion Xbb^158^ chromosome and assessing the frequency of bb^Z9^ / Xbb^158^ daughters exhibiting wild-type cuticles due to CN expansion at the bb^Z9^ locus (**Fig 2A**, see methods). We screened RNAi targeting 19 out of the 71 candidate negative regulators (represented by 28 RNAi lines) for their ability to induce ectopic rDNA magnification when expressed in the germline, based on the availability of RNAi reagents **(Table S3)**. We found that an RNAi targeting *Insulin-like receptor* (*InR*) elicited the strongest induction of rDNA magnification (RNAi line DGRC #31037), leading to 34.5% of magnified offspring **(Table S2, Fig 2B)**. This is in stark contrast to flies with normal rDNA copy number, where magnified offspring are rarely observed. Expression of a *InR^K1409A^*, a dominant negative isoform of InR, in the germline of normal rDNA CN animals (*bb^Z9^ / Ybb^+^*; *nos>InR^K1409A^*) also robustly induced rDNA magnification, confirming that reduced activity of InR induces rDNA magnification **(Fig 2A)**.

**Figure 2.**
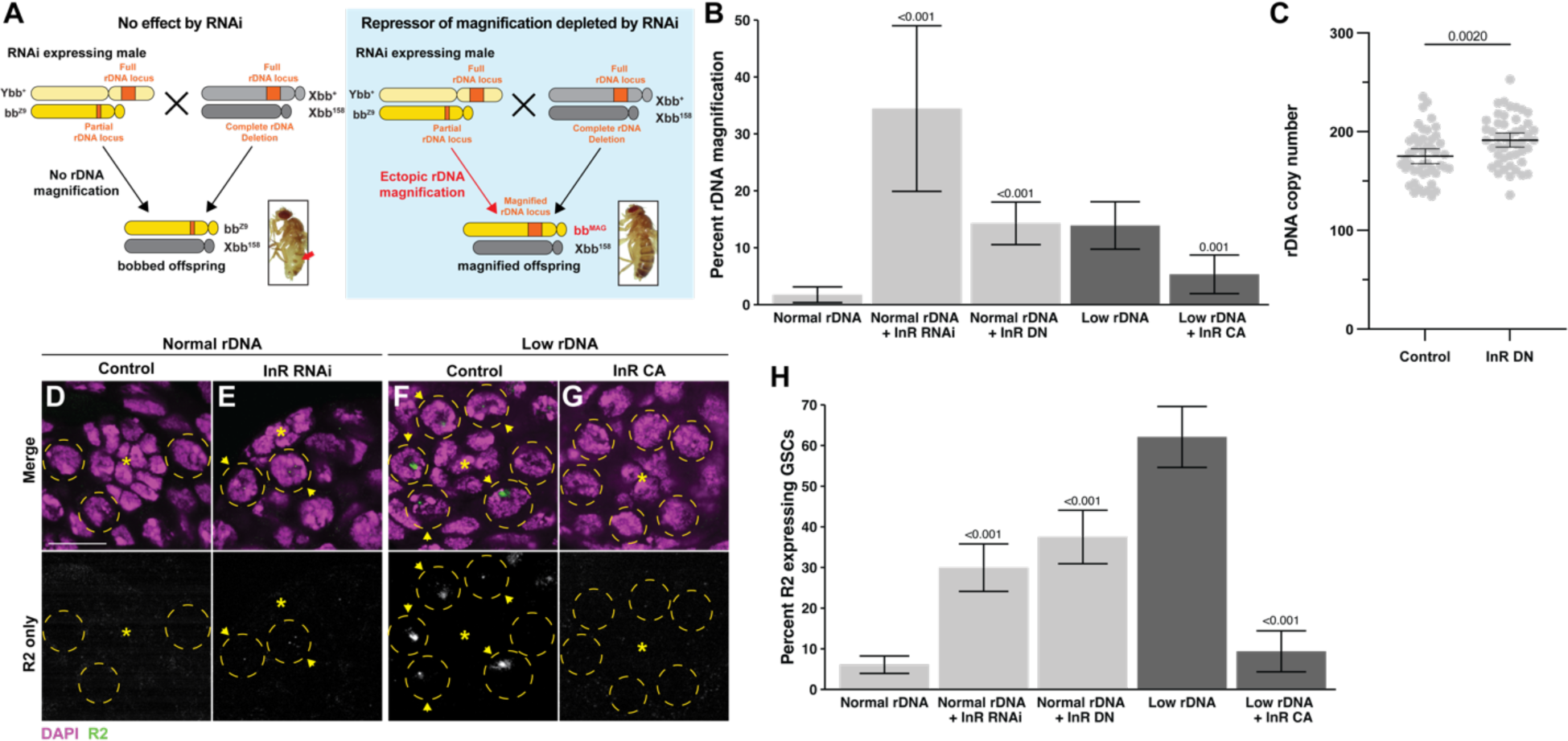
InR represses rDNA magnification and R2 expression. **A** Schematic to rapidly assess function of candidate rDNA magnification repressors. RNAi targeting a candidate gene is expressed in the early germline of males harboring the *bb^Z9^* X chromosome with Y chromosome containing a wild type rDNA locus (Ybb^+^). Males are mated to females harboring a X chromosome completely lacking rDNA (*Xbb^158^*), and the resultant *bb^Z9^ / Xbb^158^* daughters are assessed for rDNA magnification based on reversion of the *bobbed* phenotype (indicated by red arrowhead). Inset images of bobbed and reversion phenotypes from (Nelson et al, 2023b). **B** Frequency of offspring with ectopic rDNA magnification determined by presence of wild-type cuticle. P-value determined by chi-squared test compared to non-transgene condition of same rDNA content. Error: 95% CI. **C** rDNA CN assessed by ddPCR in individual *bb^Z9^ / Xbb^158^* daughters of control of InR dominant negative expressing males. N = 48 for both samples. P-value determined by Student’s t-test. Error: 95% CI. **D-G** R2 RNA FISH images in normal **(D-E)** or low rDNA CN testes **(F-G)**. DAPI in magenta, R2 in green. Asterisk (*) indicates the hub (stem cell niche), GSCs in yellow dotted circle. R2 positive GSCs indicated by yellow arrowhead. Scale bar: 10 µm. **H** Frequency of GSCs expressing R2. P-value determined by chi-squared test compared to non-transgene condition of same rDNA content. Error: 95% CI.

Moreover, quantification of rDNA copy number by droplet digital PCR (ddPCR) revealed that *bb^Z9^* chromosomes inherited from *bb^Z9^ / Ybb^+^*; *nos>InR^K1409A^* fathers have on average 16.3 more rDNA copies than *bb^Z9^* chromosomes inherited from control males (*bb^Z9^ / Ybb^+^*; *nos:Gal4* alone) (191.4 in the offspring from *nos>InR^K1409A^*father vs. 175.1 in the offspring from control father, regardless of their cuticle phenotype; p < 0.01, **Fig 2C**). Moreover, expression of a constitutively active *InR* (*nos>InR^K414P^*) substantially reduced the frequency of rDNA magnification in animals with low rDNA CN, which would usually induce strong magnification **(Fig 2B)**. Since InR is the sole receptor for IIS in Drosophila (insulin/insulin-like growth factor signaling) (Nässel et al. 2015), these results raise the possibility that rDNA magnification is controlled by cells’ major growth control pathway.

### Expression of the rDNA-specific retrotransposon R2 is repressed by InR

Expression of the rDNA-specific retrotransposon R2 in GSCs is required for rDNA magnification (Nelson et al. 2023b). R2 activity creates DNA breaks at rDNA loci that can be used for USCE to cause rDNA magnification **(Fig 1A)** (Yang et al. 1999). To determine if the effect of InR to suppress rDNA magnification is via repression of R2 expression, we used RNA FISH to examine whether downregulation of InR leads to increased R2 expression in GSCs. While R2 is typically silent in GSCs with normal rDNA CN, we indeed found that inhibition of *InR* via either RNAi (DGRC #31037) or expression of the dominant negative *InR^K1409A^*isoform increased the portion of GSCs expressing R2, despite having normal rDNA CN **(Fig 2D-H)**. Conversely, while R2 is typically frequently expressed in GSCs with low rDNA CN, expression of constitutively active *InR^K1409A^* reduced the portion of R2 expressing GSCs in this context **(Fig 2H)**. These results indicate that InR represses R2 and downregulation of InR in GSCs with low rDNA CN may allow for R2’s expression, which in turn induces rDNA magnification.

### The mechanistic Target of Rapamycin Complex 1 (mTORC1) functions downstream of InR to regulate R2 expression and rDNA magnification

There are multiple different effectors of IIS downstream of InR that can each impact transcriptional, translational, and metabolic activity (Teleman 2009), thus we sought to identify which effector(s) function in InR-mediated repression of R2 and rDNA magnification. One of the major transcriptional effectors of IIS is the transcription factor FoxO, which is phosphorylated through the Pi3K / AKT pathway upon IIS activation to sequester FoxO in the cytoplasm and prevent transcription of FoxO targets (Puig et al. 2003). We tested whether FoxO transcriptional activity may promote R2 expression and rDNA magnification. However, we found that over-expression of FoxO in GSCs did not induce ectopic rDNA magnification in animals with normal rDNA CN, and inhibition of FoxO in GSCs through RNAi did not reduce rDNA magnification in animals with low rDNA CN **(Fig S3A)**. We additionally found no difference in FoxO nuclear localization between low and normal rDNA CN GSCs **(Fig S3B-C)**. These results suggest that FoxO is not involved in rDNA magnification downstream of InR. We next tested if Pi3K / AKT activity is reduced when rDNA CN is low. We expressed a GFP-tagged pleckstrin homology domain (tGPH), which binds phosphatidylinositol-3,4,5-P_3_ (PIP_3_) (Britton et al. 2002). PIP_3_ is formed at the lipid membrane by Pi3K during InR stimulation to activate the AKT pathway, and thus membrane tGPH localization indicates active Pi3K / AKT signaling (Britton et al. 2002). We found no difference in GSC tGPH membrane localization in magnifying compared to non-magnifying conditions **(Fig S3D-E)**, indicating Pi3K activity is not reduced during rDNA magnification. Furthermore, we found that RNAi-mediated inhibition of the Pdk1 kinase, the major mediator of PIP_3_ that activates AKT (Toker and Newton 2000), does not induce ectopic rDNA magnification **(Fig S3A)**. Together, these results indicate that neither FoxO-dependent nor-independent Pi3K /AKT activity functions to repress rDNA magnification.

Having established that PI3K/Akt is unlikely the mediator of repressing rDNA magnification downstream of IIS, we tested the involvement of mTOR. mTOR is a downstream effector of IIS that regulates transcriptional activity and can be stimulated in both InR-dependent and -independent manner (Grewal 2009). We found that mTOR inhibition through rapamycin feeding induced ectopic rDNA magnification in males with normal rDNA CN **(Fig 3A)**.

**Figure 3.**
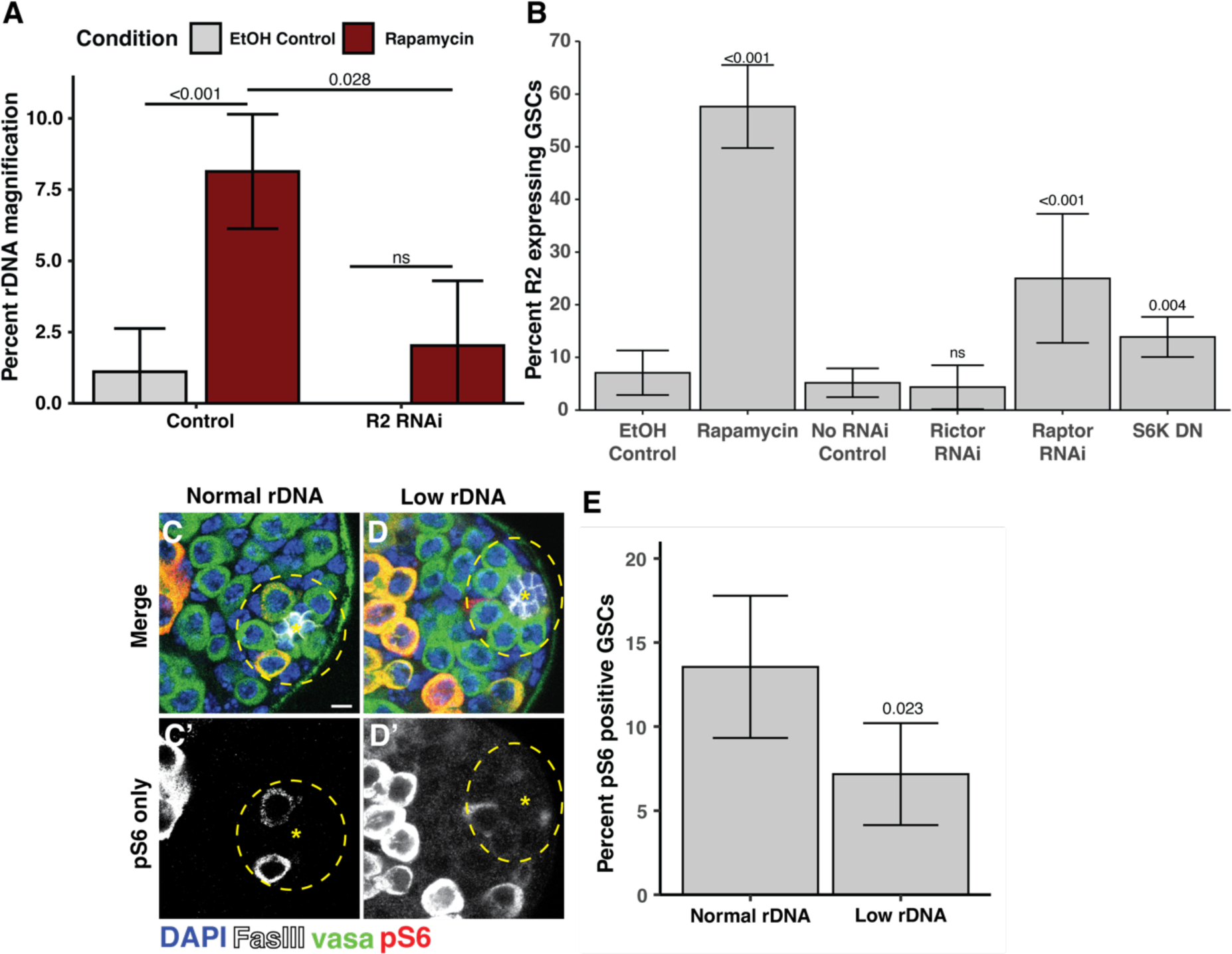
mTORC1 suppresses R2 expression and rDNA magnification. **A** Percent wild-type offspring in rDNA magnification assay from rapamycin fed or control EtOH fed males. P-values determined by chi-squared test. Error: 95% CI. **B** Percent R2 positive GSCs in rapamycin fed or EtOH fed control males and inhibition of mTor factors. P-value determined by chi-squared test. Error: 95% CI. **C-D** Images of phosphorylated S6 (pS6) in normal and low rDNA CN testes. **G** Percent pS6 positive GSCs in animals with normal or low rDNA CN. P-value determined by chi-squared test. Error: 95% CI.

Rapamycin feeding similarly increased the portion of R2 expressing GSCs in normal rDNA CN males **(Fig 3B)**. In addition, RNAi-mediated knockdown of R2 suppressed rDNA magnification caused by rapamycin **(Fig 3A)**, indicating that mTOR represses rDNA magnification via the suppression of R2. mTor functions in two complexes, mTORC1 and mTORC2 (Frappaolo and Giansanti 2023). We found that RNAi-mediated inhibition of the mTORC1-specific factor Raptor increased the portion of R2 expressing GSCs, but there was no effect of inhibiting the mTORC2-specific factor Rictor **(Fig 3B)**. We found that expression of a dominant negative isoform of S6 kinase (*S6k^K109Q^*), a major target of mTORC1 (Barcelo and Stewart 2002), also increased the portion of R2 expressing GSCs **(Fig 3D)**. Furthermore, we found that the portion of GSCs with phosphorylated ribosomal protein S6, a readout of mTORC1 activity (Kim et al. 2017; Romero-Pozuelo et al. 2017), is reduced in animals with low rDNA CN **(Fig 3D-F)** indicating that mTORC1 activity is reduced in GSCs in response to low rDNA CN. Together these results indicate that mTORC1 represses R2 expression when rDNA CN is abundant, but mTORC1 activity is reduced when rDNA CN is low, allowing for R2 expression and the induction of rDNA magnification.

### Dietary condition influences germline R2 expression and rDNA magnification activity

Because IIS and mTor are central mediators of nutrient signaling (Teleman 2009), we postulated that rDNA magnification might be influenced by nutrient conditions via these pathways. IIS and mTor stimulation by high caloric diets has been shown to reduce inherited rDNA CN (Aldrich and Maggert 2015) and we reasoned that nutrient conditions may dynamically alter rDNA magnification activity. To test this notion, we examined if changes in dietary conditions influenced R2 expression similar to modulation of InR and mTor activity. Males were fed for 5 days on varying diets containing the same amount of dietary sugar (5% sucrose), but with modified nutritional yeast (1%, 5%, and 30%). We respectively call these diets SY1, SY5, and SY30, with the SY5 diet most closely matching the nutritional content of the standard food used in our other experiments. We found that, while animals with normal rDNA CN fed on standard food (and SY5 food) rarely express R2 (only 5% of GSCs), the same animals fed on SY1 food had an increased frequency of R2 expressing GSCs **(Fig 4A-B, E)**. Conversely, animals with low rDNA CN on SY5 food exhibit a high frequency of R2-expresing GSCs, but feeding them on SY30 diets reduced the frequency in R2 expressing GSCs **(Fig 4C-E)**. These results indicate that R2 expression in GSCs is regulated in response to nutritional inputs, in a manner that induces R2 under lower nutrient conditions but suppresses its expression under high nutrient conditions.

**Figure 4.**
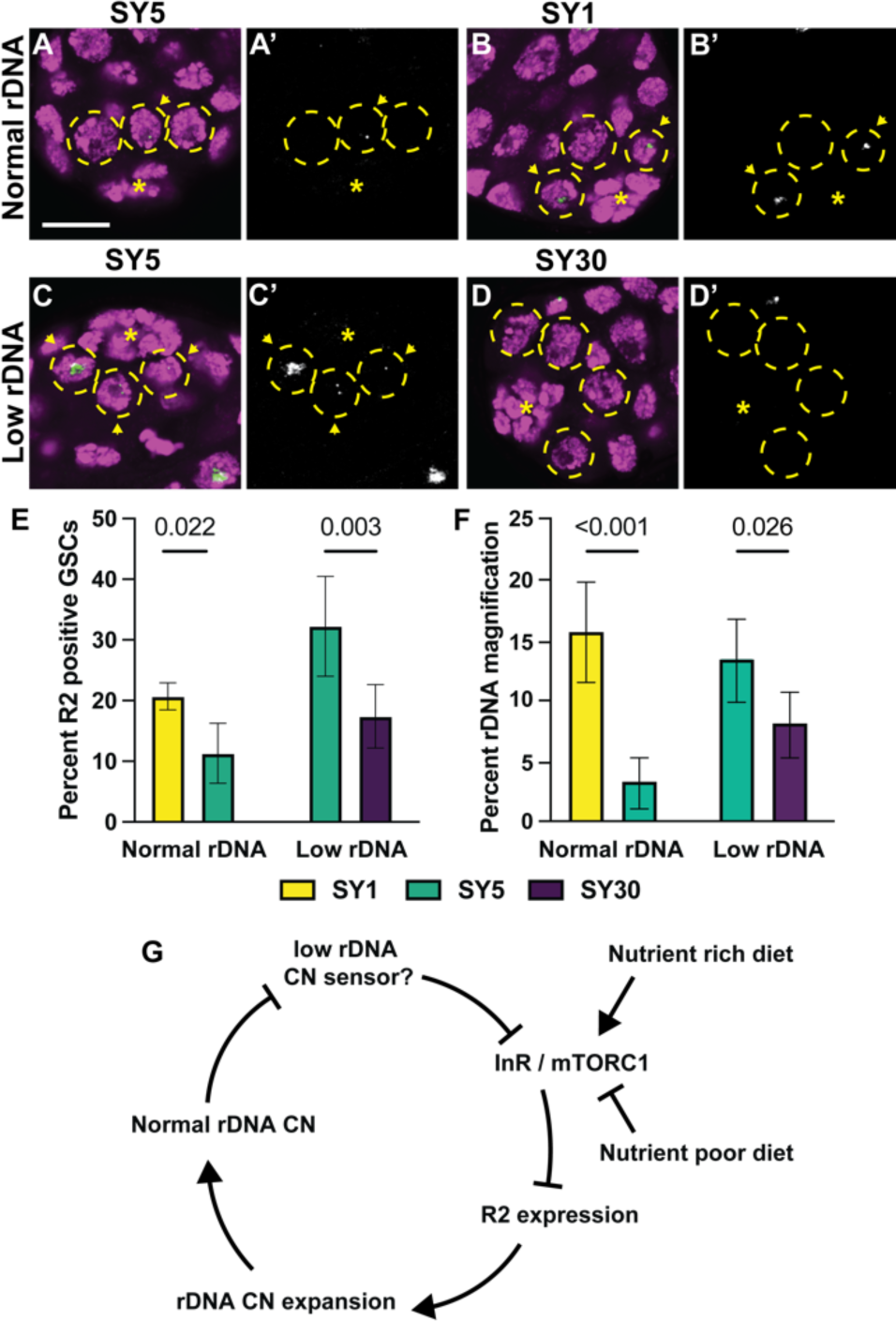
Dietary conditions alter germline R2 expression and rDNA magnification activity. **A-D** Images of R2 FISH in testes of low and normal rDNA CN. DAPI in magenta and R2 in green. **A’ – D’** are R2 channel only. **E** Percent R2 positive GSCs in males with low or normal rDNA CN on low and high calorie diets. **F** Percent offspring with wild-type cuticles in rDNA magnification assay of from males with normal or low rDNA CN fed on low and high calorie diets. Error = 95% CI. P-value determined by chi-squared test. **G** Model of dietary impact on rDNA copy number regulation through InR. InR represses R2 expression to suppress rDNA CN expansion. When rDNA CN is reduced, InR expression and mTORC1 activity is repressed via an unknown mechanism. Nutrient rich and poor diets can respectively activate or repress InR and mTORC1 activity to collectively influence R2 expression and rDNA CN expansion with rDNA CN sensing.

Moreover, we found that these dietary conditions influence rDNA CN expansion activity, suggesting that R2 expression controlled by dietary conditions is functionally linked to rDNA CN changes. Normal rDNA CN males were raised on SY1 or SY5 media for their first 10 days of adulthood, then mated to females on standard food to assess the frequency of rDNA magnification among their offspring (See methods, **Fig S4)**. We observed that rDNA magnification was very rare from males raised on SY5 media, but that the frequency of rDNA magnification was dramatically increased in males raised on SY1 media **(Fig 4F)**. On the contrary, animals with low rDNA CN, which would normally induce magnification, had reduced rDNA magnification, when fed with SY30 food for 10 days **(Fig 4F)**, indicating that this high caloric diet suppresses rDNA magnification. The low rDNA CN males raised on SY30 food still experienced some rDNA magnification, indicating that high caloric diets dampen, but do not completely suppress, rDNA magnification activity. Together, these findings reveal that dietary inputs influence R2 regulation, and in turn rDNA magnification activity, likely through impacting the IIS / mTOR pathways.

## Discussion

rDNA loci are essential but inherently unstable genetic elements, and their maintenance is particularly critical in the germline lineage for their continuation through generations. We previously showed that the rDNA-specific retrotransposon R2 plays a critical role in inducing rDNA magnification in Drosophila GSCs, representing a rare example of host-TE mutualism (Nelson et al. 2023b), instead of them being ‘genomic parasites’ as generally regarded. However, for such a mutualistic host-TE relationship to work, TEs’ activity must be precisely regulated to limit TE expression when most beneficial, while preventing unnecessary transposition that would threaten host genome integrity. That is, R2’s expression/activity must be integrated within the host’s signaling network to allow a mutually beneficial relationship.

In this study, we investigated such a mechanism by which R2 regulation is integrated in the host’s physiological system to sense rDNA CN. We used scRNA-seq to identify host factors that are differentially expressed in GSCs during rDNA magnification. Through this approach, we found that InR is a major negative regulator of rDNA magnification. *InR* transcripts were downregulated in response to low rDNA CN, and this downregulation was necessary to induce R2 expression and rDNA magnification. Moreover, we found that mTORC1 is the downstream effector of InR regulating R2 expression: mTORC1 is required to silence R2 expression when the animals have sufficient rDNA CN, whereas mTORC1 activity is downregulated in GSCs with low rDNA CN, which leads to R2 derepression and rDNA magnification. These results reveal that regulation over rDNA CN expansion and R2 expression is deeply integrated with the host’s major nutrients sensing pathway **(Fig 4G)**.

Intriguingly, we found that dietary conditions, which influence IIS and mTORC1 activity, can alter R2 expression and rDNA magnification activity. These results have an important implication: not only that animals (and GSCs within) have the ability to sense rDNA CN to regulate rDNA magnification, but that the number of rDNA copies may also be under the influence of nutrient conditions. That is, even when animals have low rDNA CN that would normally induce magnification, high nutrients dampened rDNA magnification. On the contrary, low nutrient conditions induced rDNA magnification even if animals had normal rDNA CN. It remains unclear why nutrient availability influences rDNA CN. It is possible that dietary conditions alter the physiologically required number of rDNA copies or potentially distort the physiological readout of rDNA CN. Regardless of the source of diet-induced rDNA CN changes, these findings highlight a counterintuitive relationship between nutrients and rDNA maintenance: rDNA CN is not maintained in conditions when rRNA synthesis is most needed, but rDNA CN is increased when biosynthetic resources are limited. Although counterintuitive, this aligns with the known relationship between nutrients and rDNA CN. For example, high-nutrient diets have been shown to cause somatic and germline rDNA CN reduction in Drosophila in an IIS- and mTor-dependent manner (Aldrich and Maggert 2015). Furthermore, increased mTor activity was shown to cause rDNA CN reduction in mouse hematopoetic stem cells, and some cancer types with high mTor activity are associated with low rDNA CN (Xu et al. 2017). It remains unclear as to why these conditions with high biosynthetic activity are associated with reduced rDNA CN that may hinder ribosome biogenesis. It was proposed that reduced rDNA CN may be selected for in highly mitotic cells to help expedite DNA replication, particularly in conditions of replication stress (Xu et al. 2017; Salim et al. 2017). GSC proliferation rate is associated with nutrient availability (McLeod et al. 2010), suggesting IIS and mTor may similarly optimize rDNA CN for replicative demand in GSCs. Alternatively, these observations may indicate that the mechanisms of rDNA CN expansion are not compatible with rRNA transcription, preventing rDNA CN expansion when rRNA synthesis is most active. Indeed, replication-transcription conflict is a source of CN destabilizing DNA breaks at rDNA loci in many organisms (Salim and Gerton 2019; Goffová and Fajkus 2021), suggesting similar conflict between replication and magnification may also destabilize rDNA CN. Counter to our observations in Drosophila, rDNA CN expansion in yeast is actually stimulated by mTor and suppressed by low calorie conditions (Jack et al. 2015). The differences between yeast and flies that underlie the opposing effects of nutrition on rDNA CN regulation are remain unclear. Further investigation into the specific mechanisms of rDNA CN regulation, as well as the specific physiological consequences of high and low rDNA CN in multicellular organisms, is needed to understand why rDNA CN and nutrient status sensing is integrated to dynamically regulate rDNA CN expansion.

How IIS and mTor regulate R2 expression to control rDNA magnification awaits future investigation. R2 lacks its own promoter, therefore its expression is entirely dependent on read-through transcription of the rDNA copy where it is inserted (George and Eickbush 1999). mTORC1 is a known positive regulator of rDNA transcription via its interaction with Pol I recruitment factors (Grewal et al. 2007; Ghosh et al. 2014), yet counterintuitively our data suggests that mTORC1 activity suppresses transcription at R2 containing rDNA copies. It is possible that mTORC1 differentially regulates R2-inserted vs -uninserted rDNA copies, promoting R2-uninserted rDNA copies, while negatively regulating R2-inserted rDNA copies. Indeed, mTor activity is known to increase total rRNA transcription without increasing R2 expression (Aldrich and Maggert 2015). Further investigation into the mechanisms of rDNA transcriptional regulation and the effects of mTORC1 activity on transcription of R2-containing rDNA copies is needed to fully understand how R2 is regulated in response to rDNA CN. Interestingly, IIS and mTor activity are also well described regulators of lifespan across eukaryotes, and their inhibition by dietary or genetic manipulation extends lifespan (Fontana et al. 2010), although the mechanisms that mediate these effects remain unclear (Kennedy and Lamming 2016). The instability of rDNA has been proposed to be a major factor contributing to replicative senescence in yeast (Ganley and Kobayashi 2014), and yeast mutants that increase or reduce rDNA CN stability respectively lengthen and shorten replicative lifespan (Defossez et al. 1999; Kaeberlein et al. 1999; Takeuchi et al. 2003). The possibility that IIS and mTOR inhibition may allow for rDNA CN expansion during aging suggests that rDNA CN preservation may be a mechanism for their inhibition to extend lifespan.

The nature of the mechanism that senses rDNA CN in the Drosophila germline has remained elusive since rDNA magnification was first discovered over 50 years ago. Interestingly, rDNA magnification is only active when rDNA is reduced at the Y-chromosome locus specifically(Tartof 1973), and it is more active in animals with more severe bobbed defects (Hawley and Marcus 1989), suggesting it is regulated by a combination of locus-specific features and ribosomal deficiency. Conditions that disrupt ribosome biogenesis or rRNA synthesis have been shown to lead to increased R2 expression (He et al. 2015; Fefelova et al. 2022), suggesting reduced ribosome biogenesis or function may signal cellular rDNA insufficiency. It remains unclear if ribosome biogenesis or function is actually disrupted in the germline of bobbed animals: however, we found over a quarter of genes encoding for ribosomal proteins have reduced expression in GSCs with low rDNA CN **(Table S1**, 22.9-fold enrichment, p<10^-15^ determined by chi-squared test). This enrichment for ribosomal proteins to have reduced expression implies that ribosome activity is indeed reduced in low rDNA CN condition and may serve as a sensor of reduced rDNA CN, though these effects may be the indirect result of reduced mTor activity. The involvement of IIS and mTOR in regulating rDNA magnification suggests low rDNA CN may be sensed by these pathways through changes in metabolic conditions, such as glucose or amino acid availability (Teleman 2009). Indeed, many ribosome deficiencies are associated with altered glycolysis, amino acid synthesis, and proteosome activity (Kampen et al. 2019), suggesting metabolism may become altered when rDNA CN is reduced. Our finding that dietary manipulation impacts R2 expression and rDNA CN expansion activity further suggests a key role for metabolite availability in rDNA CN sensing. Alternatively, stabilized p53 is a common consequence of disrupted ribosome biogenesis, including in Drosophila female GSCs (Chakraborty et al. 2011; Martin et al. 2022), which may repress insulin and mTor activity in GSCs with reduced rDNA CN. Further investigation into how rDNA CN is sensed by the cell and the potential role of metabolites in this function is critical to understanding the physiological consequences of the integrated sensing of environmental and genomic conditions to regulate rDNA CN.

Taken together, the present work demonstrates that repression of the rDNA-specific retrotransposon R2 by the IIS / mTor pathway regulates rDNA maintenance activity in GSCs. We propose that this role for IIS in rDNA regulation integrates nutrient and rDNA CN regulation to transgenerationally maintain and adjust rDNA CN. Such a dynamic mechanism may explain the widespread intra-species variation in rDNA CN observed in many organisms (Klootwijk et al. 1979; Dawid et al. 1978; Wellauer and Dawid 1974; Smirnov et al. 2021). Excessive mTor activation has been associated with some instances of rDNA CN reduction in mice and humans (Xu et al. 2017), suggesting its function to regulate rDNA CN may be widely conserved across animals. These observations may reflect a conserved nature of rDNA CN dynamics and regulation to integrate natural CN fluctuation with changing environmental factors such as nutrients.

## Methods

### Single cell RNA sequencing sample preparation

50 testes were hand dissected from 1 to 5-day-old flies in 1x PBS and transferred immediately into tubes with cold 1x PBS on ice. Tissue dissociation was performed as previous described (Raz et al. 2023). Cell viability and density was determined on a hemocytometer using Trypan Blue stain and DIC imaging. Cells were processed for library preparation using the 10 X Genomics Chromium Controller and Chromium Single Cell Library and Gel Bead Kit following standard manufacturer’s protocol. Amplified cDNA libraries were quantified by bioanalyzer, size selected by AMPure beads, and sequenced on a NovaSeq SP.

### Single cell sequencing data analysis

The 10 X CellRanger pipeline was used with default parameters to map reads to the DM3 reference genome. The resulting matrices were read and processed into a single combined data set using the Seurat R package. Cells containing between 5,500 and 250,000 counts and 200 and 9000 features were isolated for data analysis to enrich for analysis of intact isolated cells. The relative similarity and differences in gene expression for all 20,138 combined cells from low rDNA CN and normal rDNA CN nos>upd samples was reduced using 14 dimensions and a resolution of 0.5 in a Uniform Manifold Approximation and Projection (UMAP)-based dimensionality reduction for cluster analysis. Previously determined expression signatures for distinct spermatogenesis developmental stages and testis cell types were used to initially assign cell type designations to the UMAP clusters (Raz et al. 2023). Specifically, clusters that predominantly contained *vas*, *nos*, and *ovo* expressing cells were considered GSC & spermatogonia (SG); *dlg1*, *CadN*, *tj*, and *zfh1* labeled Cyst cell clusters; *fzo* and *CycB* labelled spermatocytes; and *CG32106* and *m-cup* identified spermatids. Any unlabeled clusters were subsequently categorized based on the cell type associated with their strongest unique expression markers in previous analysis (Raz et al. 2023).

GSCs were determined for differential expression analysis based on sub-clustering and expression selection methods. Sub-clustering renormalized data from cells in GSC / SG containing clusters and a Cyst cell cluster outgroup and gene expression differences were again reduced using 14 dimensions for UMAP-based dimensionality reduction. GSCs containing clusters were distinguished from SG based on the high concentration of cells expressing *nos*, *vas*, and *ovo*, while SG containing clusters were *vas* positive, but mostly devoid of *nos* and *ovo* expression. Cyst cell containing clusters were again identified by *dlg1*, *CadN*, *tj*, and *zfh1*, and clusters containing both GSC and Cyst cell makers were considered a mixed population cluster and excluded from analysis. For expression selection methods, cells from any original cluster were isolated based on positive expression for *vas, nos,* and *ovo*, but negative expression for *tj* and *zfh1*. Genes with a log_2_ fold-change greater than 0.25 and a Bonferroni corrected p-value less than 0.05 were considered significantly differentially expressed. Potential false-positive expression increases due contaminating ambient RNA from lysed cells were eliminated by removing all genes with increased expression both low rDNA CN somatic cells and GSCs.

### DNA isolation and rDNA copy number measurement by droplet digital PCR (ddPCR)

DNA was isolated from individual Drosophila adults using a modified DNeasy Blood and Tissue DNA extraction kit (Qiagen), as previously described (Nelson et al. 2023b). All DNA samples were quantified and purity was determined by NanoDrop One spectrophotometer (ThermoFisher). rDNA copy number quantification was determined by ddPCR by assessing 28S copy number normalized to control genes (RpL49 and Upf1) as previously described (Nelson et al. 2023b). Primers and probes are listed in **Table S4**. ddPCR droplets were generated from samples using QX200 Droplet Generator (Bio-Rad) with ddPCR Supermix for Probes (No dUTP) (Bio-Rad). PCR cycling was completed on a C100 deep-well thermocycler (Bio-Rad) and fluorescence was measured by the QX200 Droplet Reader (Bio-Rad) and calculated using Quantasoft software (Bio-Rad).

### rDNA magnification assay

For rDNA magnification assays, *Ybb^0^* / *bb^Z9^*males were used for ‘low rDNA’ conditions and Y / *bb^Z9^* males were used for ‘normal rDNA’ conditions. To assess the frequency of rDNA magnification, males were mated in bulk to *bb^158^* / *FM6, Bar* females. Resulting *bb^Z9^ / bb^158^* female offspring were selected based on the absence of the *Bar* dominant marker and scored for the presence of the *bobbed* phenotype. The portion of offspring having wild-type cuticles and not the *bobbed* phenotype represents the frequency of rDNA magnification.

### RNA FISH

RNA FISH samples were prepared using an R2 Stellaris probe set (Biosearch Technologies) as previously described (Lu et al. 2018). Dissected testes were fixed for 30 min in 4% formaldehyde in PBS, briefly washed twice in PBS for 5 minutes, and permeabilized in 70% ethanol at 4°C overnight. Samples were briefly washed in 2x saline-sodium citrate (SSC) with 10% formamide, then hybridized with 50 nM probes at 37°C overnight. Following hybridization, samples were washed twice with 2x SSC containing 10% formamide for 30 min and mounted in VECTASHIELD with DAPI (Vector Labs). Samples were imaged using a Leica Stellaris 8 confocal microscope with a 63x oil-immersion objective and processed using Fiji (ImageJ) software.

### Immunofluorescence

Immunofluorescence staining of testes was performed as previously described (Cheng et al. 2008). Testes were dissected in 1x PBS and fixed in 4% formaldehyde in PBS for 30 min. Following fixing, testes were briefly washed two times in 1x PBS containing 0.1% Triton-X (PBS-T), followed by a 30-minute wash in PBS-T. Samples were subsequently incubated at 4°C overnight with primary antibody in 3% bovine serum albumin (BSA) in PBS-T. Samples were washed three times for 20 minutes in PBS-T, then incubated with secondary antibody in 3% BSA in PBS-T at 4°C overnight. Samples were then washed three times again for 20 min in PBS-T and mounted in VECTASHIELD with DAPI (Vector Labs). Primary antibodies used in this study can be found in **Table S4**. Images were taken with a Leica Stellaris 8 confocal microscope with 63x oil-immersion objectives and processed using Fiji (ImageJ) software.

### Drosophila genetics and dietary conditions

Drosophila lines used in this study can be found in **Table S4**. All animals were reared at 25°C on standard Bloomington medium, except when specific diets are mentioned. Experimental diets consisted of 1% agar, 5% sucrose, and respectively 1%, 5%, or 30% yeast for SY1, SY5, and SY30 diets. Animals were raised on standard medium, and newly eclosed adult males were transferred to experimental diets for 5 days prior to dissections for RNA FISH, or 10 days prior to magnification assay.

### Rapamycin feeding

10 µM Rapamycin food was prepared by adding Rapamycin dissolved in ethanol directly to vials containing standard corn-meal based food. Control food was prepared by adding an equal volume to ethanol. 15 newly eclosed adult males were placed in vials and raised at 25°C for 5 days. After the feeding course, males were mated to females on standard food for magnification assays.

### Statistics

Statistical significance was determined by chi-squared test for all comparisons of the percentage of samples with categorical values (percent magnification or R2 positive cells), and error bars were generated using the Confidence Interval for a Population Proportion formula. All comparisons of samples with independent values (rDNA copy number), significance was determined by Student’s t-test, and error represents 95% confidence interval. Significance for differential gene expression analysis was determined by a non-parametric Wilcoxon rank sum test. All represented p-values are Bonferroni corrected based on the number of comparisons.

## Acknowledgements

We thank the Bloomington Drosophila Stock Center, Kyoto Drosophila Stock Center, FlyBase, and Developmental Studies Hybridoma Bank for reagents and resources. We thank Dr. Aurelio Teleman and Dr. Jongkyeong Chung for sharing reagents. We thank the Yamashita lab members for discussion and comments on the manuscript. We also thank the Whitehead Institute Genome Technology Core for their assistance with scRNA-seq. This research was supported by NIH F32GM143850 (A.A.R), the Howard Hughes Medical Institute (Y.M.Y.) and the John Templeton Foundation (Y.M.Y).

## Author Contributions

JON: Conceptualization, Design, Data acquisition, Data analysis, Data curation, Writing – original draft, Writing – review & editing, Data visualization. AS: Data acquisition, Data analysis, Data curation, Writing – review & editing. AAR: Data acquisition, Data analysis, Writing – original draft, Writing – review & editing, Funding acquisition. YMY: Conceptualization, Design, Data analysis, Writing – original draft, Writing – review & editing, Data visualization, Project administration, Funding acquisition.

